# Pan-cancer survival modeling reveals structural limits of genomic feature integration in immunotherapy outcomes

**DOI:** 10.64898/2026.04.15.718634

**Authors:** Hassan Wasiu Oluwagbenga, Sola Adeleke

## Abstract

**Background:** Immune checkpoint inhibitors (ICIs) have improved outcomes across multiple cancer types, yet reliable predictors of survival remain limited. While genomic features such as tumor mutational burden (TMB) are widely used, their contribution to predictive modeling in heterogeneous real-world cohorts remains unclear. We evaluated the relative contributions of clinical and whole-genome sequencing (WGS) features in pan-cancer survival modeling.

**Methods:** We analyzed 658 patients treated with ICIs with matched WGS data from the Genomics England. Using a leakage-controlled machine learning framework with strict train–test separation, we compared four models: TMB-only, clinical-only, clinical+TMB, and an integrated 11-feature clinico-genomic XGBoost survival model. Model performance was assessed using Harrell’s concordance index (C-index) with bootstrap confidence intervals.

**Results:** TMB alone demonstrated near-random discrimination (C-index 0.50; 95% CI 0.44–0.56). Clinical variables substantially improved predictive performance (0.59; 95% CI 0.53–0.64), with marginal gain from adding TMB (0.59). The integrated model achieved a C-index of 0.60 (95% CI 0.55–0.65). While improvement over TMB alone was significant, incremental gain beyond optimized clinical models was modest. Feature attribution analysis showed that model performance was dominated by clinical variables, with genomic features contributing limited additional signal.

**Conclusions:** These findings suggest that, in heterogeneous pan-cancer cohorts, predictive performance is constrained by the underlying data structure, in which dominant clinical signals overshadow genome-scale features. This study highlights fundamental limitations in integrating genomic data into survival models across diverse cancer types and provides a benchmark for future computational approaches.

## 1. Background

Immune checkpoint inhibitors (ICIs) targeting programmed cell death protein 1 (PD-1), programmed death ligand 1 (PD-L1), and cytotoxic T-lymphocyte–associated antigen 4 (CTLA-4) have fundamentally reshaped the treatment landscape across multiple advanced malignancies. Landmark trials demonstrated durable responses in melanoma and non-small cell lung cancer (NSCLC), establishing PD-1/PD-L1 blockade as a therapeutic cornerstone in modern oncology [1, 2]. Subsequent expansion into urothelial carcinoma, renal cell carcinoma, and head and neck cancers confirmed the broad applicability of immune checkpoint modulation. However, despite these transformative advances, a durable survival benefit is achieved in only a subset of patients. Population-level analyses estimate that the proportion of patients deriving meaningful survival benefit from ICIs remains limited relative to the total treated population [3]. Primary resistance, early progression, and immune-related toxicities continue to challenge indiscriminate immunotherapy deployment. In real-world cohorts, survival heterogeneity is even more pronounced due to advanced disease burden, prior lines of therapy, and variability in baseline performance status. Accordingly, accurate prediction of survival following ICI initiation remains one of the most pressing unmet needs in precision oncology.

Tumor mutational burden (TMB) emerged as a promising pan-cancer biomarker based on the neoantigen hypothesis, the premise that increased somatic mutation load generates immunogenic peptides recognizable by cytotoxic T cells. Early translational work demonstrated correlations between the burden of nonsynonymous mutations and response to PD-1 blockade in NSCLC [4]. Extensive pan-cancer analyses further suggested that higher TMB was associated with improved survival following immunotherapy across multiple tumor types [5]. These findings contributed to tissue-agnostic regulatory approval of TMB as a biomarker for immunotherapy eligibility. However, subsequent analyses revealed substantial variability in predictive performance across tumor contexts. Not all high-TMB tumors respond, and certain low-TMB malignancies demonstrate meaningful immunotherapy benefit. Comprehensive evaluations have shown that TMB lacks consistent predictive discrimination when applied uniformly across heterogeneous cancer populations [6]. Several biological explanations underlie this inconsistency. TMB quantifies mutation burden but not mutation quality; it does not capture clonality or neoantigen presentation efficiency; it ignores tumor-intrinsic immune evasion pathways (e.g., STK11 and KEAP1 alterations) [7]; and it fails to incorporate host clinical fitness. Moreover, reported concordance indices for TMB-only survival models in real-world cohorts are frequently modest (~0.50–0.60), reflecting limited discriminatory capacity in heterogeneous settings. Collectively, these observations suggest that while TMB may function as a coarse eligibility biomarker, it is insufficient as a standalone survival predictor in diverse pan-cancer populations. From a modeling perspective, this raises the question of whether limited performance arises from the biological inadequacy of TMB alone, or from structural challenges in learning predictive signals across heterogeneous cancer types.

Whole-genome sequencing (WGS) provides a more comprehensive representation of tumor biology than aggregate mutation counts. Beyond total mutational burden, WGS enables characterization of mutational signatures reflecting endogenous and exogenous DNA-damage processes [8], structural variants, pathway-level genomic instability, and resistance-associated driver alterations. For example, homologous recombination deficiency (HRD)-associated signatures reflect impaired double-strand break repair and have been linked to genomic instability and altered immune landscapes. Similarly, ultraviolet (UV) and tobacco-associated signatures capture carcinogenic exposures that heavily influence tumor immunogenicity. Large-scale national initiatives, including the Genomics England 100,000 Genomes Project, have enabled the integration of population-scale WGS data with longitudinal clinical outcomes, creating unprecedented opportunities to examine genome-wide predictors of immunotherapy survival. Despite this potential, translating high-dimensional WGS features into clinically actionable survival prediction remains computationally challenging. The “curse of dimensionality”, sparsity of rare variants, and cross-tumor biological heterogeneity can easily dilute gene-specific signals when aggregated across cancer types. Whether WGS-derived features provide meaningful incremental survival prediction beyond established clinical variables in real-world pan-cancer immunotherapy cohorts, therefore, remains an open question.

Machine learning (ML) approaches have been proposed as powerful tools for integrating complex clinico-genomic feature spaces [9]. Algorithms such as gradient boosting machines and random survival forests can model nonlinear interactions and higher-order feature dependencies that may be inaccessible to traditional Cox proportional hazards frameworks. However, enthusiasm for ML in oncology has been heavily tempered by methodological concerns. Large-scale methodological reviews have highlighted frequent issues, including data leakage, overfitting due to limited sample sizes, optimistic cross-validation without independent hold-out testing, and inadequate uncertainty reporting [10, 11]. In the context of WGS data, additional risks emerge. Technical sequencing artifacts, including contamination scores and tumor purity estimates, can function as hidden proxies for tumor burden. If not explicitly excluded, high-capacity ML models may exploit these artifacts, producing artificially inflated performance that collapses under independent validation. Therefore, rigorous train-test separation, careful feature curation, and bootstrap-based uncertainty estimation are essential for realistic performance assessment. In particular, the interaction between feature heterogeneity and model capacity remains poorly understood, especially when combining clinical and genomic variables in pan-cancer datasets.

Significantly, survival following ICI initiation is influenced not only by tumor-intrinsic genomics but also by systemic host factors. Baseline functional status, commonly measured using the Eastern Cooperative Oncology Group (ECOG) performance scale [12], remains one of the strongest predictors of oncology outcomes across treatment modalities. Patients with impaired performance status exhibit reduced physiologic reserve, impaired immune competence, and limited tolerance to therapy. Similarly, the number of prior systemic therapy lines reflects cumulative disease burden and therapeutic resistance. These systemic clinical determinants may dominate survival discrimination in heterogeneous real-world cohorts, potentially overshadowing incremental genomic contributions. Whether integrated ML models meaningfully exceed optimized clinical baselines or primarily repackage existing clinical signal remains a critical translational question.

In this study, we performed a rigorous real-world evaluation of integrated clinico-genomic machine learning models for survival prediction in a heterogeneous pan-cancer cohort of patients treated with immune checkpoint inhibitors. Using strict 70/30 train-test separation and 1000-iteration bootstrap uncertainty estimation, we compared tumor mutational burden alone, clinical-only Cox modeling, a clinical + TMB baseline, and an integrated 11-feature clinico-genomic XGBoost survival model. We hypothesized that baseline clinical burden would remain the dominant determinant of survival discrimination; that whole-genome–derived features would provide statistically measurable but modest incremental refinement; and that leakage-controlled modeling would yield more conservative performance estimates than those reported in cross-validated genomic ML studies. By explicitly quantifying the relative contribution of clinical and genomic features, this study aims to provide insight into the structural limits of predictive modeling in heterogeneous pan-cancer datasets.

## 2. Results

A total of 658 patients met the inclusion criteria and formed the final analytic cohort. The cohort represented a heterogeneous, real-world pan-cancer population treated with immune checkpoint inhibitors across multiple solid tumor types, predominantly driven by lung cancer, melanoma, renal cell carcinoma, and urothelial carcinoma. At baseline, the median patient age was 72.0 years. The cohort was predominantly male, comprising 56.4% males and 43.6% females. Regarding baseline functional status, 38.9% of patients had an Eastern Cooperative Oncology Group (ECOG) performance status of 0. In comparison, 48.8% had a score of ≥ 1 (with the remainder unrecorded, reflecting standard real-world registry limitations). Furthermore, patients had received a median of 0.0 prior lines of systemic therapy, indicating a predominantly first-line immunotherapy population. The median follow-up time for the entire cohort was sufficient to capture mature survival events, enabling robust time-to-event modeling. The cohort selection process and final analytic population are summarized in Figure 4.

Model discriminative performance was evaluated on the independent, locked test cohort using Harrell’s Concordance Index (C-index) with 1000-iteration bootstrap 95% confidence intervals (CIs). The baseline single-analyte model utilizing Tumor Mutational Burden (TMB) alone demonstrated limited predictive discrimination (C-index 0.5047; 95% CI: 0.4447–0.5592), performing only marginally better than random chance. In contrast, the Clinical-only baseline model, which relied exclusively on standard host factors (e.g., age, sex, performance status, line of therapy), achieved substantially higher predictive accuracy (C-index 0.5855; 95% CI: 0.5299– 0.6395). The addition of TMB to the clinical baseline provided minimal incremental benefit (C-index 0.5928; 95% CI: 0.5364–0.6482). The proposed integrated XGBoost-AFT framework, utilizing the curated 11-feature clinico-genomic space, achieved the highest overall discrimination with a C-index of 0.6015 (95% CI: 0.5495–0.6530). While this integrated approach provided a statistically significant improvement over TMB alone (p=0.006), it offered only a modest incremental lift ($\Delta$ = +0.0087) over the standard Clinical + TMB baseline. This suggests that while high-dimensional WGS features contain a measurable prognostic signal, their additive value is constrained when baseline clinical burden is adequately modeled. A comparative overview of model performance across all baseline and integrated models is presented in **Figure 1**, illustrating the marginal incremental gain achieved by the integrated clinico-genomic framework over clinical baselines.

**Figure 1:**
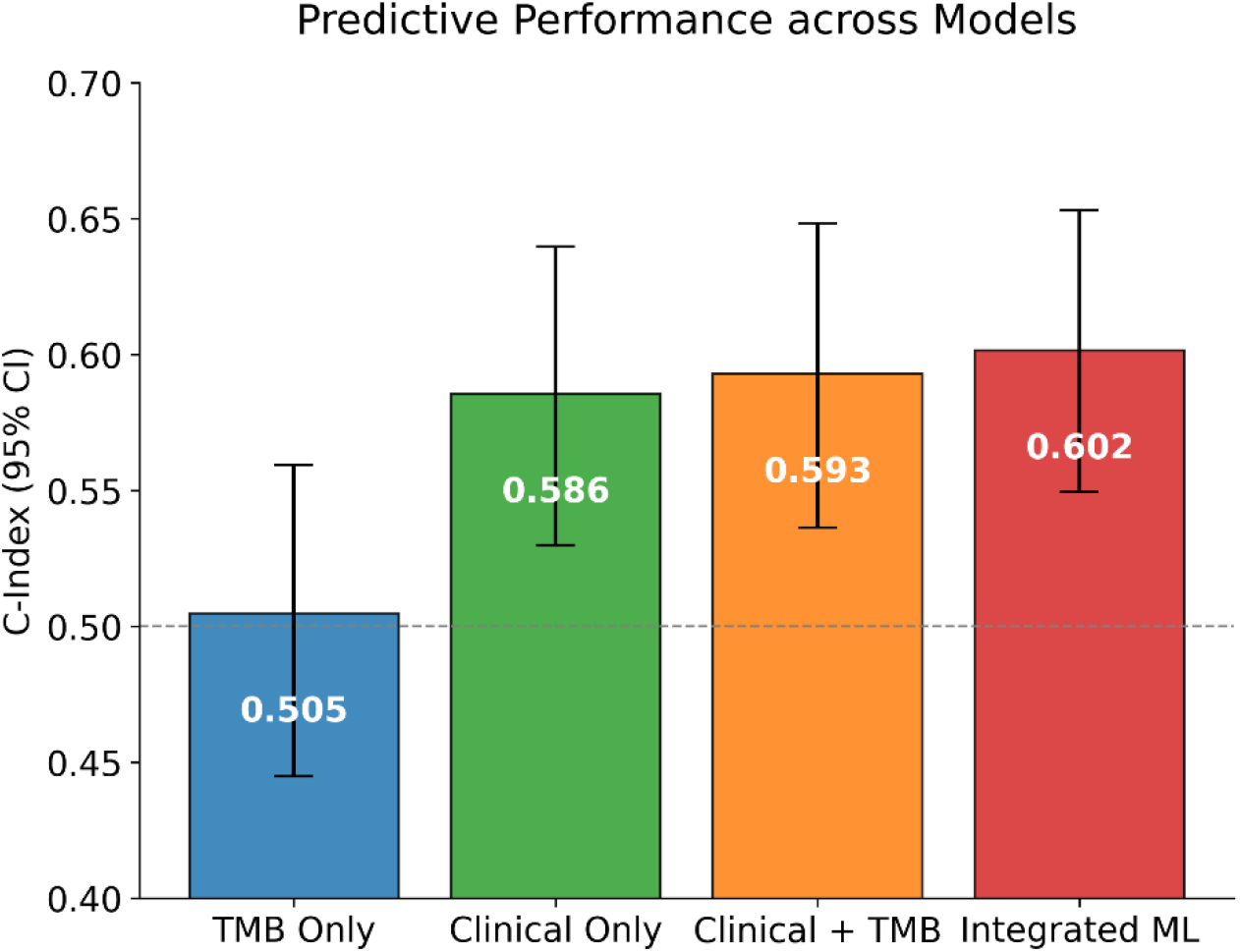
Model performance comparison (C-index with 95% CI)

To translate the continuous XGBoost-AFT output into clinically actionable strata, the test cohort was dichotomized into “High-Risk” and “Low-Risk” groups based on the median predicted survival score. To assess the model’s clinical utility, survival stratification was evaluated using Kaplan–Meier analysis. Kaplan-Meier survival analysis revealed a profound and statistically significant divergence in overall survival (OS) trajectories between the two strata. Patients classified in the High-Risk group exhibited substantially accelerated mortality compared to the Low-Risk cohort, yielding a Hazard Ratio (HR) of 1.96 (95% CI: 1.45–2.64; log-rank $p < 0.001$). The survival separation between groups is visualized in Figure 2.

**Figure 2.**
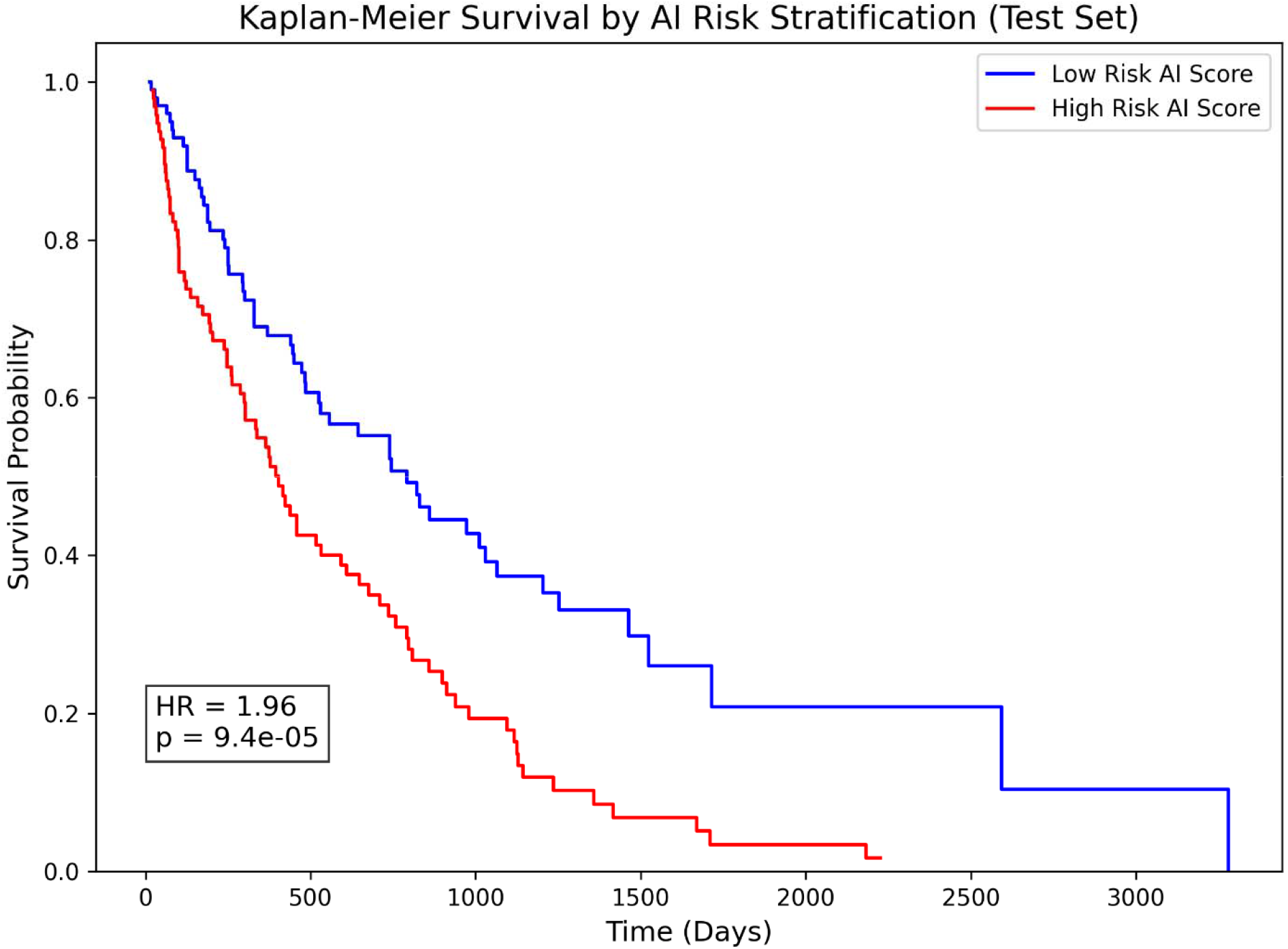
Kaplan–Meier curves for ML-defined high vs low risk

To interpret model predictions and identify key drivers of survival, SHAP-based feature importance analysis was performed (Figure 3). The SHAP summary analysis revealed that baseline ECOG Performance Status was the single most dominant determinant of survival prediction, heavily outweighing individual genomic metrics. Among the integrated whole-genome features, mutational signatures exhibited the highest predictive importance. Specifically, the Ultraviolet (UV) radiation signature and the Homologous Recombination Deficiency (HRD) signature emerged as top-tier predictors of extended survival, aligning with established biological paradigms of immune microenvironment modulation. Conversely, the model autonomously prioritized specific somatic variants, notably pathogenic alterations in *KEAP1* and *TP53*, as primary drivers of innate immunotherapy resistance and shortened survival, confirming the biological fidelity of the learned predictive pathways.

**Figure 3.**
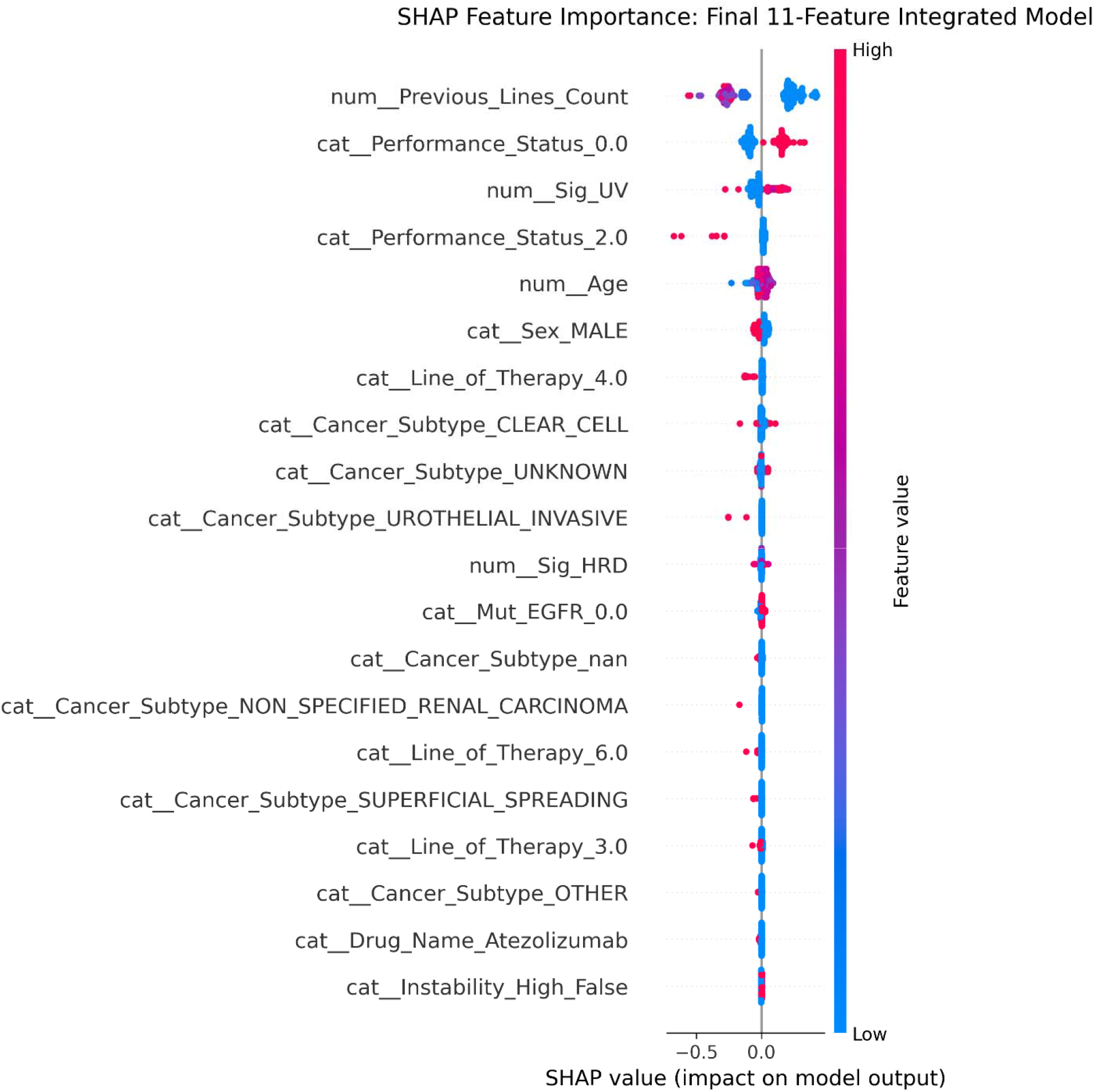
SHAP summary plot (11-feature integrated model)

**Figure 4.**
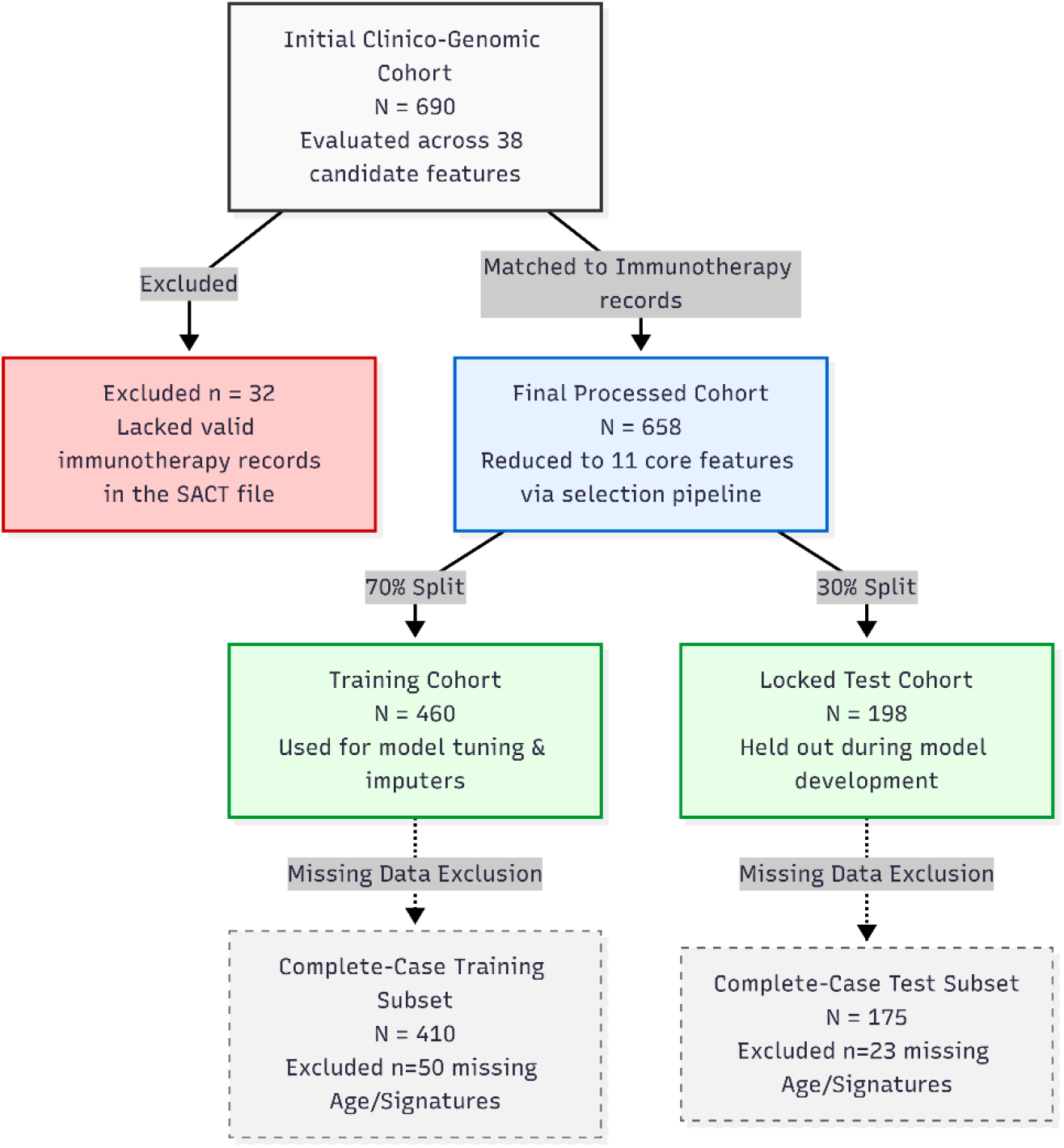
Cohort Inclusion Summary

To rigorously validate the model’s structural integrity, two prespecified sensitivity analyses were conducted. First, a complete-case analysis was performed by restricting the cohort strictly to patients with 0% missing data across all 11 features. The complete-case integrated model achieved a C-index of 0.5917, representing a negligible reduction compared to the median-imputed model ($\Delta$ = −0.0098). This minimal variance confirmed that the model’s performance was not an artifact of the imputation strategy. Second, a feature-ablation study was conducted by removing ECOG Performance Status from the integrated model. This ablation resulted in a marked decline in predictive accuracy (C-index 0.5727; $\Delta$ = −0.0288). This substantial performance drop definitively demonstrated the model’s heavy reliance on systemic clinical burden. It reinforced the conclusion that genome-scale ML models must benchmark against robust clinical baselines to avoid overstating the independent prognostic value of genomic data.

**Table 1:** Sensitivity and ablation performance summary.

| Model / Analysis Configuration | Concordance Index (C-index) | 95% CI | $\Delta$ from Main Model | Description |
| --- | --- | --- | --- | --- |
| Baseline Benchmarks |  |  |  |  |
| TMB-Only | 0.5047 | 0.4447 – 0.5592 | -0.0968 | Genomic baseline using Tumor Mutational Burden alone. |
| Clinical-Only | 0.5855 | 0.5299 – 0.6395 | -0.0160 | Clinical baseline excluding all WGS features. |
| Clinical + TMB | 0.5928 | 0.5364 – 0.6482 | -0.0087 | Standard combined baseline. |
| Main Proposed Framework |  |  |  |  |
| Integrated ML (11 Features) | 0.6015 | 0.5495 – 0.6530 | Ref | Final XGBoost-AFT clinico-genomic model. |
| Sensitivity & Ablation Analyses |  |  |  |  |
| Complete-Case Analysis | 0.5917 | *Not calculated* | -0.0098 | Restricted to patients with 0% missing data (Age, UV, HRD) to rule out imputation bias. |
| Feature Ablation: Excl. Performance Status | 0.5727 | *Not calculated* | -0.0288 | Removal of the dominant clinical predictor to assess model reliance on baseline fitness. |

Abbreviations: CI, Confidence Interval; TMB, Tumor Mutational Burden; WGS, Whole Genome Sequencing; ML, Machine Learning; Ref, Reference for comparison

## 3. Discussion

These findings suggest that predictive performance in pan-cancer survival modeling is constrained by the underlying data structure, in which dominant clinical signals limit the marginal contribution of genome-scale features. First, baseline clinical burden, specifically functional performance status, remains the dominant determinant of survival, heavily outweighing individual genomic metrics. Second, Tumor Mutational Burden (TMB), when used as a standalone continuous predictor, exhibits inadequate discriminative capacity ($C$-index = 0.5047) in heterogeneous real-world populations. Third, while integrating high-dimensional Whole-Genome Sequencing (WGS) features into a gradient-boosting architecture provides a statistically significant improvement over TMB alone, the incremental prognostic lift beyond a robust clinical baseline is modest ($\Delta$ = +0.0087).

The poor predictive performance of the TMB-only model aligns with mounting clinical skepticism regarding its universal applicability. While initial retrospective analyses established TMB as a tissue-agnostic biomarker for pembrolizumab eligibility [16], subsequent multi-cohort evaluations have demonstrated that high TMB does not uniformly correlate with CD8+ T-cell infiltration or survival benefit across all histologies [6]. Furthermore, aggregate mutation counting fails to distinguish between clonal and subclonal neoantigens, the latter of which are highly inefficient at driving sustained anti-tumor immune responses [17]. Our findings underscore that in unselected, real-world pan-cancer cohorts, TMB should not be deployed as an isolated prognostic tool, as it lacks the resolution to capture tumor-intrinsic immune evasion or host-level clinical frailty.

Despite the modest aggregate lift in the $C$-index, the SHAP interpretability analysis confirmed that the XGBoost-AFT model autonomously learned highly accurate, biologically plausible oncology principles. The algorithm ranked the Ultraviolet (UV) radiation signature and Homologous Recombination Deficiency (HRD) as top-tier predictors of extended survival. This reflects known biological paradigms: UV-driven melanomas frequently exhibit highly immunogenic, clonal neoantigen landscapes, while HRD-driven genomic instability promotes cytosolic DNA accumulation, thereby activating the cGAS-STING pathway and priming the tumor microenvironment for ICI response [18]. Conversely, the model correctly identified somatic variants in *KEAP1* and *TP53* as powerful drivers of innate resistance and accelerated mortality. *KEAP1* mutations, frequently co-occurring with *STK11* in lung adenocarcinomas, are well-established mediators of “cold” tumor microenvironments characterized by T-cell exclusion and profound resistance to PD-(L)1 blockade [7]. The model’s ability to independently prioritize these specific molecular vulnerabilities validates the utility of WGS over targeted panels. From a computational perspective, these results highlight a key limitation of pan-cancer modeling: the dilution of feature-specific signal across heterogeneous disease contexts. While genomic alterations may be strongly predictive within individual tumor types, their effects are attenuated when aggregated across biologically distinct cancers, reducing their contribution to global model performance.

The ablation of ECOG performance status resulted in a severe degradation in model accuracy ($\Delta$ = −0.0288), highlighting a fundamental truth often obscured in the genomic ML literature: real-world survival is primarily dictated by systemic host fitness. Patients with impaired functional status (ECOG ≥ 1) often suffer from cancer-induced cachexia, underlying comorbidities, and an inability to tolerate immune-related adverse events (irAEs). Recent real-world analyses have consistently shown that poor baseline ECOG status drastically truncates overall survival following ICI initiation, independent of genomic biomarkers [19]. Clinical trials rigorously exclude frail patients, artificially magnifying the predictive value of genomics. In real-world data, however, the sheer physiologic burden of advanced disease imposes a “predictive ceiling” that even the most advanced multi-omic algorithms cannot bypass.

Several limitations warrant consideration. First, pan-cancer aggregation introduces immense biological heterogeneity; tumor-specific modeling frameworks might uncover stronger, histology-specific genomic signals that are currently diluted in pooled cohorts. Second, our feature space was restricted to DNA-level WGS metrics and standard clinical data. The absence of spatial transcriptomics, T-cell receptor (TCR) repertoire profiling, and multiplex immunofluorescence of the tumor microenvironment limits the model’s predictive ceiling [20]. Finally, while internally validated via a rigorous locked hold-out set within the UK NHS infrastructure, external validation across independent, international cohorts remains necessary to confirm broader generalizability.

In a highly regulated, leakage-controlled real-world pan-cancer cohort, baseline clinical burden remains the dominant determinant of survival following immune checkpoint inhibition. While whole-genome sequencing offers superior biological resolution over isolated TMB, successfully identifying critical mutational signatures and resistance variants, its integration into machine learning architectures provides only modest incremental refinement for overall survival prediction. These findings establish realistic performance expectations for AI in clinical oncology and mandate that future genomic ML frameworks benchmark heavily against robust clinical baselines to ensure actual translational utility.

## 4. Methods

### 4.1 Study Design and Cohort Selection

This retrospective cohort study used clinico-genomic data from the National Genomic Research Library (NGRL), accessed through the Genomics England Research Environment, including data generated by the Genomics England 100,000 Genomes Project [13]. We established a pan-cancer cohort comprising 658 patients with advanced solid malignancies treated with immune checkpoint inhibitors (ICIs) (e.g., pembrolizumab, nivolumab, atezolizumab). The index date was defined as the initiation of ICI therapy. The primary endpoint was overall survival (OS), defined as the time from ICI initiation to death from any cause. Patients alive at the last follow-up were right-censored at the date of last clinical contact. This study was conducted using de-identified data under an approved Genomics England research protocol. All participants provided informed consent for genomic sequencing and data linkage. The study adhered to the Declaration of Helsinki and was conducted in accordance with ethical and governance frameworks established by Genomics England and the National Health Service (NHS).

### 4.2 Genomic and Clinical Data Extraction

Tumor and matched normal WGS data were processed using the standardized GEL bioinformatics pipeline. Somatic single-nucleotide variants (SNVs), insertions/deletions (indels), and structural variants were quantified. Tumor mutational burden (TMB) was calculated as the total number of nonsynonymous somatic mutations per megabase (mut/Mb). Beyond TMB, we extracted single-base substitution (SBS) mutational signatures using the COSMIC v3.2 reference framework [8], specifically isolating signatures with established immunomodulatory roles, including the Ultraviolet (UV) radiation signature and the Homologous Recombination Deficiency (HRD) signature. Targeted somatic alterations in known drivers of ICI resistance (e.g., *KEAP1, TP53, EGFR*) were also annotated [7]. Baseline clinical variables extracted from the longitudinal registry included patient age at diagnosis, sex, specific cancer subtype, Eastern Cooperative Oncology Group (ECOG) performance status [12], and the number of prior lines of systemic therapy.

### 4.3 Feature Engineering and Artifact Exclusion

The initial dataset comprised 38 raw clinical and multi-omic variables. To prevent high-dimensional overfitting and ensure strict translational utility, a rigorous feature selection pipeline was applied prior to final model training. First, variables exhibiting severe missingness (>80%, e.g., particular rare mutational signatures) were excluded. Second, to prevent data leakage and artificial inflation of model performance, a prevalent methodological flaw in genomic ML [10], technical WGS artifacts, specifically sequencing Contamination_Score and Tumour_Purity, were explicitly removed. Exploratory analyses demonstrated that high-capacity algorithms could exploit these technical metrics as artificial proxies for tumor burden and sample quality. Finally, empirical feature importance evaluation (via recursive feature elimination) was utilized to exclude noisy, non-contributory variables that degraded pan-cancer generalizability. This dimensionality-reduction pipeline yielded a highly curated final feature space comprising 11 core clinico-genomic predictors. All feature preprocessing, including encoding, scaling, and imputation, was performed using only the training data, with parameters applied to the test set to prevent data leakage.

### 4.4 Handling of Missing Data

Missingness was assessed exclusively among the 11 features retained in the final integrated model. Eight of the eleven variables exhibited complete data (0% missingness). Three variables (Age, HRD Signature, UV Signature) demonstrated moderate missingness (11.09% each). To preserve sample size while avoiding leakage, missing values were imputed within the training set using median imputation for continuous variables, and the fitted imputer was subsequently applied to the locked test set without refitting [11]. To rigorously evaluate the potential impact of imputation on model performance, a prespecified complete-case sensitivity analysis was conducted, restricting both the training and test cohorts to participants with no missing values across all 11 features. Missingness was assessed prior to model development, and imputation was performed within the training set only to avoid information leakage.

### 4.5 Machine Learning Pipeline and Survival Modeling

The total cohort (N=658) was partitioned using a strict 70/30 train-test split, yielding a training set (N=460) and an independent, locked test set (N=198). To capture complex, non-linear clinico-genomic interactions, Hyperparameter tuning was conducted using cross-validation within the training set only. Model selection was finalized prior to evaluation on the test set. We deployed an eXtreme Gradient Boosting Accelerated Failure Time (XGBoost-AFT) survival architecture [14]. Unlike standard Cox proportional hazards models, the AFT framework directly models the expected survival time, providing enhanced flexibility for heterogeneous pan-cancer datasets. Hyperparameter tuning (e.g., learning rate, maximum tree depth, L1/L2 regularization) was strictly confined to the training set using 5-fold cross-validation to optimize the concordant index and prevent overfitting.

### 4.6 Statistical Analysis and Model Interpretability

Model discriminative performance was evaluated on the locked test set using Harrell’s Concordance Index (C-index). To estimate the uncertainty of the performance metrics, 95% confidence intervals (CIs) were generated using a 1000-iteration bootstrap resampling approach on the test set. For baseline comparison, three benchmark models were constructed: a TMB-only model, a Clinical-only model, and a combined Clinical + TMB model. Risk stratification was assessed by dichotomizing the test cohort into “High-Risk” and “Low-Risk” groups based on the median predicted survival time, evaluated via Kaplan-Meier survival curves and the log-rank test. Statistical significance for Δ C-index comparisons was assessed using bootstrap resampling, with two-sided p-values derived from the empirical distribution. Finally, to overcome the “black-box” nature of the XGBoost algorithm, Shapley Additive exPlanations (SHAP) were employed [15]. SHAP values provided unified, biologically plausible feature importance rankings, quantifying the precise predictive contribution of each clinical and genomic variable for each patient. All analyses were conducted using Python (v3.11). The overall data processing pipeline and machine learning framework are illustrated in Figure 5.

**Figure 5.**
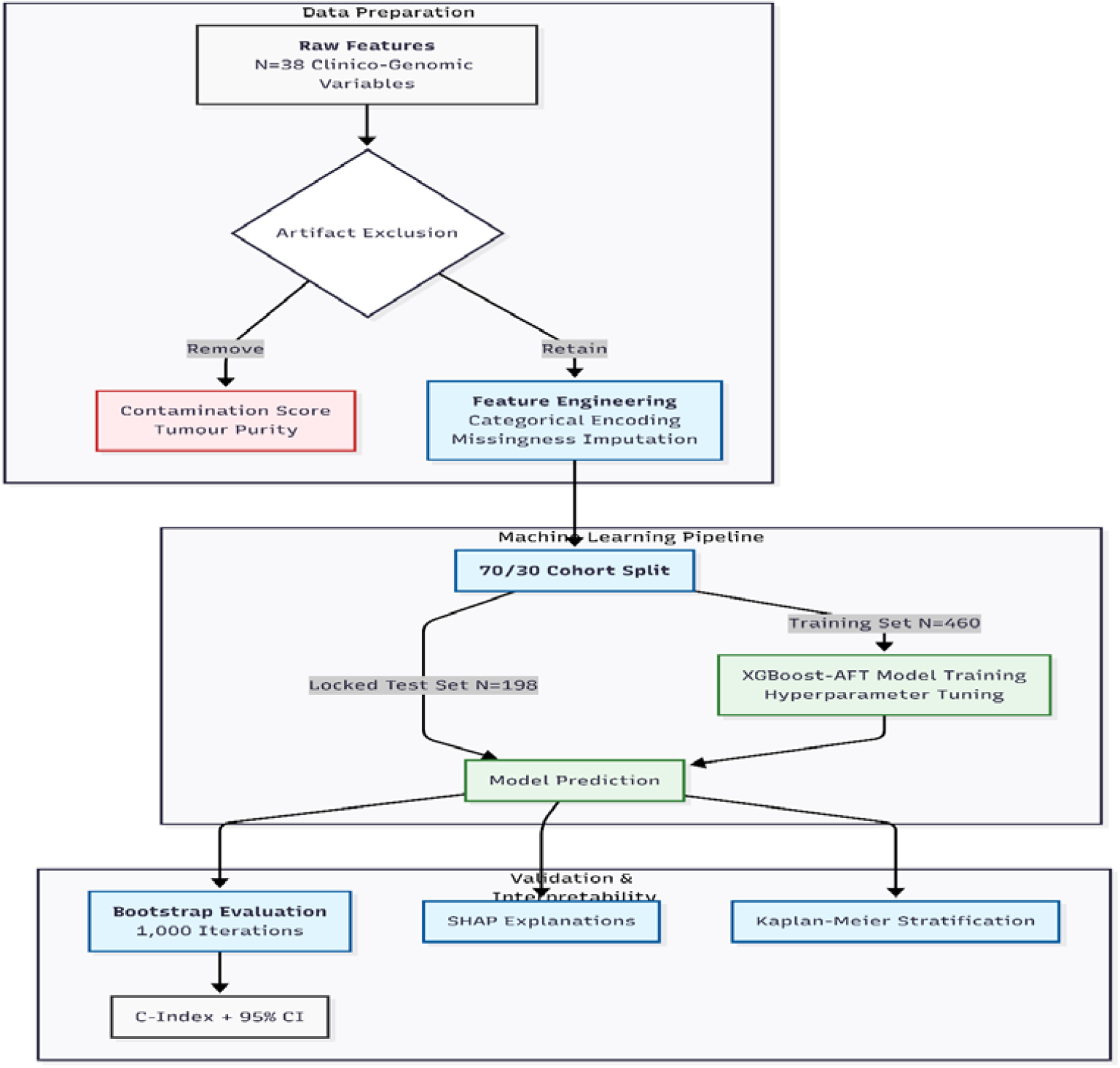
Data preparation and ML framework

### 4.7 Data Availability

Data from the National Genomic Research Library (NGRL) used in this research are available within the secure Genomics England Research Environment [21]. Access to NGRL data is restricted to adhere to consent requirements and protect participant privacy.

Data used in this study include whole-genome sequencing–derived variant calls, mutational signatures, and associated clinical variables for participants treated with immune checkpoint inhibitors within the Genomics England research environment. These datasets were accessed and analyzed within the secure research infrastructure and include derived datasets generated for survival modeling and machine learning analyses within the project workspace. Cohort construction and feature extraction were performed using curated datasets within the research environment, including cancer registry data, SACT treatment data, genome file path mappings, and cancer pathology datasets.

Access to NGRL data is provided to approved researchers who are members of the Genomics England Research Network, subject to institutional access agreements and research project approval under participant-led governance. For more information on data access, visit: https://www.genomicsengland.co.uk/research. Code used for analysis is available upon reasonable request Through the Secure Genomics England AWS working environment

## Acknowledgements

We gratefully acknowledge the participants of the National Genomic Research Library (NGRL), whose contributions made this research possible. Secure access to the NGRL under project ID RR1390 was provided by Genomics England, which delivers the NGRL in partnership with NHS England, and is wholly owned by the UK Department of Health and Social Care.

The NGRL contains participants’ health data collected by the NHS as part of their care, along with samples and data from their participation in research, for which fully informed consent has been obtained. This includes genomic and clinical data provided through the NHS Genomic Medicine Service, as well as data obtained through research studies, including the 100,000 Genomes Project and the Generation Study, both of which are delivered in partnership with the NHS, and from other research cohorts involving external collaborators.

